# Oncogenic Mutations in Histologically Normal Endometrium: The New Normal?

**DOI:** 10.1101/561050

**Authors:** Vivian Lac, Tayyebeh M. Nazeran, Basile Tessier-Cloutier, Rosalia Aguirre-Hernandez, Arianne Albert, Amy Lum, Jaswinder Khattra, Teresa Praetorius, Madeline Mason, Derek Chiu, Martin Köbel, Paul J. Yong, Blake C. Gilks, Michael S. Anglesio, David G. Huntsman

## Abstract

The presence of somatic driver mutations in endometriosis has previously been believed to represent early events in transformation, however our group and others have described such mutations in roughly one-third of cases of deep infiltrating or iatrogenic endometriosis. These forms of endometriosis rarely progress to malignancy. Recent studies have also shown somatic driver mutations in normal skin, blood, peritoneal washings, and esophageal epithelium. Such findings prompt speculation on whether such mutations exist in the eutopic endometrium – the likely tissue of origin of endometriosis. In the current study we investigated the presence of somatic driver mutations in histologically normal endometrium from women lacking evidence of gynecologic malignancy or endometrial hyperplasia. Twenty-five women who underwent hysterectomies and 85 women who underwent endometrial biopsies were included in this study. Formalin-fixed, paraffin-embedded tissue specimens were analyzed by means of targeted sequencing followed by orthogonal validation with droplet digital PCR. PTEN and ARID1A immunohistochemistry (IHC) was also performed as surrogates for inactivating mutations in the respective genes. Overall, we observed somatic driver-like events in over 50% of histologically normal endometrial samples analyzed, which included hotspot mutations in *KRAS*, *PIK3CA*, and *FGFR2* as well as PTEN-loss by IHC. Analysis of anterior and posterior samplings collected from women who underwent hysterectomies is consistent with the presence of somatic driver mutations within clonal pockets spread throughout the uterus. The prevalence of such oncogenic mutations also increased with age (OR: 1.05 (95% CI: 1.00 – 1.10), p = 0.035). These findings have implications on our understanding of aging and so-called “normal tissues”, thereby necessitating caution in the utilization of mutation-based early detection tools for endometrial or other cancers.

## Introduction

Endometriosis is a chronic disease affecting approximately 10 percent of reproductive-aged women. It leads to a variety of clinical symptoms including chronic (often cyclical) pain, dyspareunia, dysmenorrhea, and infertility[1,2]. Endometriosis is the major risk factor for clear cell and endometrioid carcinomas of the ovary (endometriosis-associated ovarian cancers; EAOCs) [3–5]. Observations of endometriotic lesions (consisting of endometrial-like glands and stroma) contiguous with EAOCs combined with molecular findings from several sequencing studies have further established endometriosis as the precursor of EAOCs[6–8]. In these studies, oncogenic mutations in genes such as *ARID1A* and *PIK3CA*[7,8] have been traced back to endometriotic lesions within the same patients, thereby implicating such alterations as early events towards malignant transformation. However, we have found somatic driver mutations in approximately one-third of deep infiltrating endometriosis and iatrogenic endometriosis cases (forms of endometriosis with little malignant potential)[9,10]. These findings suggest that such mutations may be an inherent feature of endometriosis itself, perhaps conferring survival advantages that allow lesions to persist ectopically. However, in order to study this hypothesis, it is important to clarify whether uterine epithelial mutations can pre-exist endometriosis and be present in the cells prior to implantation at an ectopic site.

In the current study, we sought to explore the mutational landscape of the eutopic (uterine) endometrium from women lacking evidence of gynecologic malignancy by means of targeted sequencing and immunohistochemistry (IHC) studies. Although the origins of endometriosis remain contentious, the concept of retrograde menstruation stands as a leading theory. Herein, endometrial fragments (originating from the uterus) are refluxed upwards through the fallopian tubes and are implanted within the pelvic cavity[11]. This theory is supported by several lines of evidence including the high prevalence of endometriosis in females with congenital outflow obstruction[12] and the successful generation of endometriotic lesions in primate models of endometriosis via intrapelvic injection of menstrual endometrium[13] or cervical occlusion[14]. Considering the dynamic nature of the endometrium as it undergoes many cycles of regeneration and tissue breakdown throughout a woman’s reproductive years[15] as well as the recent description of somatic mutations in normal tissues including the skin[16], blood[17], peritoneal washings[18], and the esophageal epithelium[19], it is plausible that somatic mutations may pre-exist in the eutopic endometrium. Moreover, a series of previous studies have described “latent precancers” (alterations in otherwise normal tissue with unknown contribution to long-term cancer risk) in the form of PTEN-null and PAX2-null glands within histologically normal endometrium[20–22]. Application of next-generation sequencing techniques will broaden our understanding of latent precancers, or pre-existing somatic alterations, in the eutopic endometrium. This is particularly important since these tissues often serves as a reference for molecular studies on endometriosis, endometrial cancer, and EAOCs.

## Materials and Methods

### Patient Identification and Specimen Collection

We obtained formalin-fixed and paraffin-embedded (FFPE) endometrial tissue specimens from 25 women who underwent hysterectomies (Supplementary Table S1) and 85 women who underwent either dilation and curettage (D&C) or endometrial biopsies (Supplementary Table S2). As confirmed by pathologist review, all women lacked evidence of gynecologic malignancy or endometrial hyperplasia and all tissue specimens were obtained from the Department of Anatomical Pathology at the Vancouver General Hospital in Vancouver, Canada. For hysterectomy patients (Hx cohort), we collected and subsequently analyzed two blocks to represent endometrial tissue from the posterior uterus and anterior uterus from each patient. For D&C or endometrial biopsy patients (Bx cohort), we targeted collection of endometrial tissue specimens to represent women of various ages (ranging from 20 – 61 years of age). Specimen, data collection and experiments were approved by the UBC BC Cancer Agency Research Ethics Board [H05-60119, H02-61375 and H08-01411].

### Sample Processing and DNA Extraction

Hysterectomy specimens were sectioned at 8µm onto glass slides, deparaffinized with xylene and stained with 10% diluted hematoxylin and eosin (H&E). Specimens were manually macrodissected to enrich the epithelial layer using the tip of a 20-guage needle under a stereo microscope. DNA from hysterectomy specimens was extracted using the ARCTURUS® PicoPure® DNA Extraction Kit (ThermoFisher Scientific, USA). D&C and endometrial biopsy specimens were cut into 3 – 6 10µm tissue scrolls. DNA from D&C and endometrial biopsy specimens was extracted using the QIAamp® DNA FFPE Tissue Kit (QIAGEN, USA). All DNA was quantitated using the Qubit 2.0 Fluorometer (ThermoFisher Scientific, USA).

### Targeted Sequencing

All specimens were sequenced using FIND IT^™^ version 3.4 (Contextual Genomics, Canada), a proprietary hypersensitive cancer hotspot assay which covers over 120 hotspots and 17 exons spanning 33 genes (Supplementary Table S3)[23]. Libraries were constructed using 75ng of total DNA input. Candidate somatic variants for orthogonal validation were selected from defined cancer hotspots reported in the Catalogue of Somatic Mutations in Cancer (COSMIC)[24], with probability scores ≥ 0.8 (indicating the likelihood that a variant belongs to the mutation class as opposed to the artifact class) and variant allele frequency (VAF) ≥ 0.8%.

### Orthogonal Validation via ddPCR

Droplet digital polymerase-chain-reaction (ddPCR) was used to orthogonally validate hotspot mutations. DNA from each specimen was pre-amplified for 10 cycles before subsequent droplet generation using the QX200 Droplet Generator (Bio-Rad Laboratories, USA)[25]. In most cases, the same aliquot of extracted DNA used for targeted sequencing was also used for ddPCR validation (Supplementary Table S5 and S6). After thermal cycling, the QX200 Droplet Reader (Bio-Rad Laboratories, USA) was used to quantify droplets. See Supplementary Table S7 for a list of primers used and Supplementary Table S8 for ddPCR assay conditions.

### PTEN and ARID1A Immunohistochemistry

Loss of PTEN immunoreactivity was used as a surrogate for PTEN loss-of-function mutations. Following the protocol outlined in our previous study[10], specimens were stained on the Ventana Discovery Ultra (Ventana Medical Systems, USA) immunostainer using a 1:25 dilution of rabbit monoclonal antibody, 138G6 (Cell Signaling, USA). Similarly, loss of nuclear ARID1A immunoreactivity was used as a surrogate for ARID1A loss-of-function mutations[10,26,27]. Specimens were stained on the Dako Omnis (Agilent Technologies, USA) automated immunostainer using a 1:2000 dilution of ARID1A rabbit monoclonal antibody EPR13501 (ab182560, Abcam, USA). TMN and BT-C. scored all ARID1A and PTEN immunostained slides.

## Results

### Somatic Driver Mutation Frequencies in Normal Endometrium

As shown in Figure 1, we found that 16 of 25 women (64%) who underwent hysterectomy and 43 of 85 women (51%) who underwent D&C or endometrial biopsy harbored at least one somatic alteration within their endometrial tissue samplings. Given our sample sizes, the prevalence of these alterations (whether specifically or overall) did not significantly differ between Hx and Bx cohorts (Supplementary Table S9). The most common somatic alterations were hotspot *KRAS* and *PIK3CA* mutations (affecting 31 of 110 and 14 of 110 total patients respectively), and focal loss of PTEN expression (defined by absence of staining in a small cluster of glands among normally expressing glands) (30 of 110 total patients) (Supplementary Figure S1). No patients exhibited loss of ARID1A either focally or globally.

**Figure 1.**
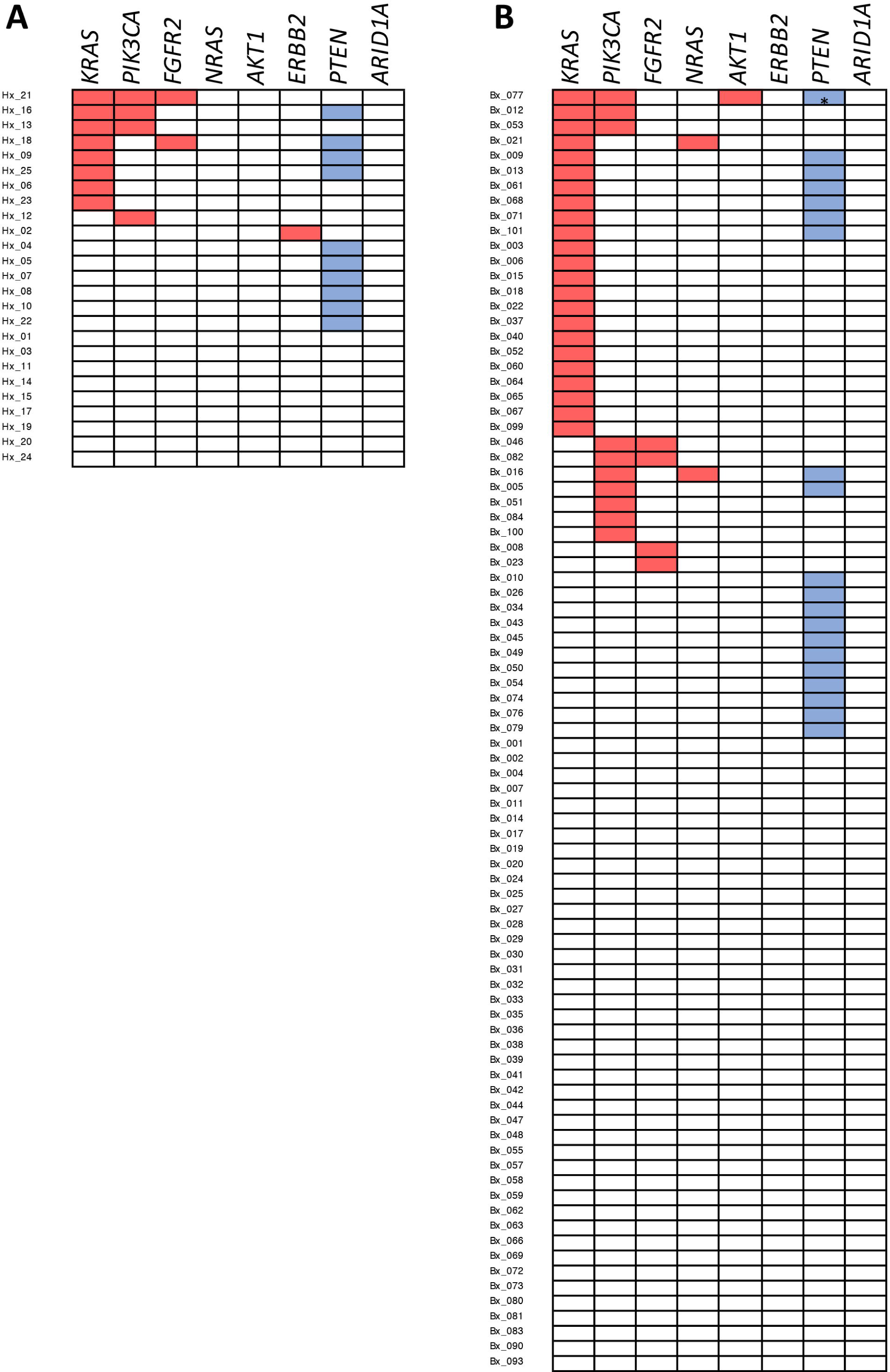
Overview of somatic, oncogenic alterations detected in the eutopic endometrium from (A) hysterectomy patients or (B) dilation and curettage or endometrial biopsy patients. Red rectangles denote driver mutations detected by targeted sequencing, whereas blue rectangles denote (zonal) loss detected through immunohistochemistry studies. The asterisk (*) denotes a patient wherein zonal loss of PTEN as well as a *PTEN* driver mutation was detected.

### Anatomical Distribution of Oncogenic Changes

For each woman in the Hx cohort, we independently collected and sequenced DNA from two tissue blocks representing endometrial tissue from the anterior and posterior uterus within the same patient. Somatic hotspot mutations were either completely absent (15 of 25 patients) or dissimilar (9 of 25 patients) between anterior and posterior samplings (Supplementary Figure S2). A single patient (Hx_25) harbored a *KRAS* G12A mutation in both samplings, however anatomical distortion of the uterus in this patient resulted in an inability to determine the spatial distinction between the two blocks of endometrial tissue (Table 1).

**Table 1.** Mutational findings within normal endometrium samples from hysterectomy patients and corresponding variant allele frequencies (VAF) determine through targeted sequencing and <u>droplet digital</u> PCR (ddPCR).

| Patient and Block ID | Region of Uterus Sampled | PTEN IHC | Driver Mutation Identified | VAF (%) - Sequencing | VAF (%) - ddPCR |
| --- | --- | --- | --- | --- | --- |
| Hx_2A | anterior | normal |  |  |  |
| Hx_2B | posterior | normal | ERBB2 S310F | 0.974 | 0.792 |
| Hx_4A | anterior | loss - zonal |  |  |  |
| Hx_4B | posterior | normal |  |  |  |
| Hx_5A | anterior | loss - zonal |  |  |  |
| Hx_5B | posterior | normal |  |  |  |
| Hx_6A | anterior | normal | KRAS G12V | 2.654 | 1.63 |
| Hx_6B | posterior | normal | KRAS G12A | 1.408 | 1.08 |
| Hx_7A | posterior | loss - zonal |  |  |  |
| Hx_7B | anterior | normal |  |  |  |
| Hx_8A | posterior | loss - zonal |  |  |  |
| Hx_8B | anterior | loss - zonal |  |  |  |
| Hx_9A | anterior | normal |  |  |  |
| Hx_9B | posterior | loss - zonal | KRAS G13D | 2.16 | 1.95 |
| Hx_10A | n/a | normal |  |  |  |
| Hx_10B | n/a | loss - zonal |  |  |  |
| Hx_12A | anterior | normal | PIK3CA R88Q | 1.337 | 1.23 |
| Hx_12B | posterior | normal |  |  | - |
| Hx_13A | posterior | normal | PIK3CA H1047R | 2.497 | 3.36 |
|  |  |  | KRAS G12C | 1.036 | 0.538 |
| Hx_13B | anterior | normal |  |  |  |
| Hx_16A | anterior | normal | KRAS G12V | 1.079 | 1.06 |
|  |  |  | PIK3CA E542K | 1.015 | 0.897 |
| Hx_16B | posterior | loss - zonal | PIK3CA H1047R | 2.402 | 1.92 |
| Hx_18A | anterior | normal | FGFR2 K659E | 5.505 | 7.77 |
|  |  |  | KRAS G12V | 1.551 | 1.24 |
| Hx_18B | posterior | loss - zonal |  |  |  |
| Hx_21A | anterior | normal | KRAS G12D | 5.587 | 4.71 |
|  |  |  | PIK3CA M1043I | 2.39 | 2.27 |
| Hx_21B | posterior | normal | KRAS G12V | 6.112 | 6.32 |
|  |  |  | PIK3CA H1047R | 2.373 | 2.76 |
|  |  |  | FGFR2 S252W | 1.889 | 1.17 |
| Hx_22A | anterior | loss - zonal |  |  |  |
| Hx_22B | posterior | normal |  |  |  |
| Hx_23A | anterior | n/a |  |  |  |
| Hx_23B | posterior | normal | KRAS G12D | 11.866 | 10.2 |
| Hx_25A | n/a | normal | KRAS G12A | 1.584 | 1.60 |
| Hx_25B | n/a | loss - zonal | KRAS G12A | 1.670 | 1.00 |

\*Data for nine patients without detectable somatic mutations or abnormal immunohistochemistry findings are omitted from this table.

### Driver Mutations Across Age

Accumulation of somatic alteration with increasing age has previously been suggested even in the context of normal tissues[16–19]. To assess the relationship between age and presence of somatic cancer-driver alterations, we generated a logistic regression model based on the sequencing and IHC findings for the 25 women in the Hx cohort and 85 women in the Bx cohort (Figure 2). Based on our model, the likelihood of a woman harbouring a somatic alteration in her endometrial tissue increases by 5% per year (OR = 1.05, 95% CI = 1.00 – 1.10, p = 0.035, Wald test). Moreover, a model incorporating both age and menstrual phase (proliferative versus secretory phase) indicates that the prevalence of mutation is independent of menstrual phase (p = 0.6309, Wald test) (Supplementary Figure S3).

**Figure 2.**
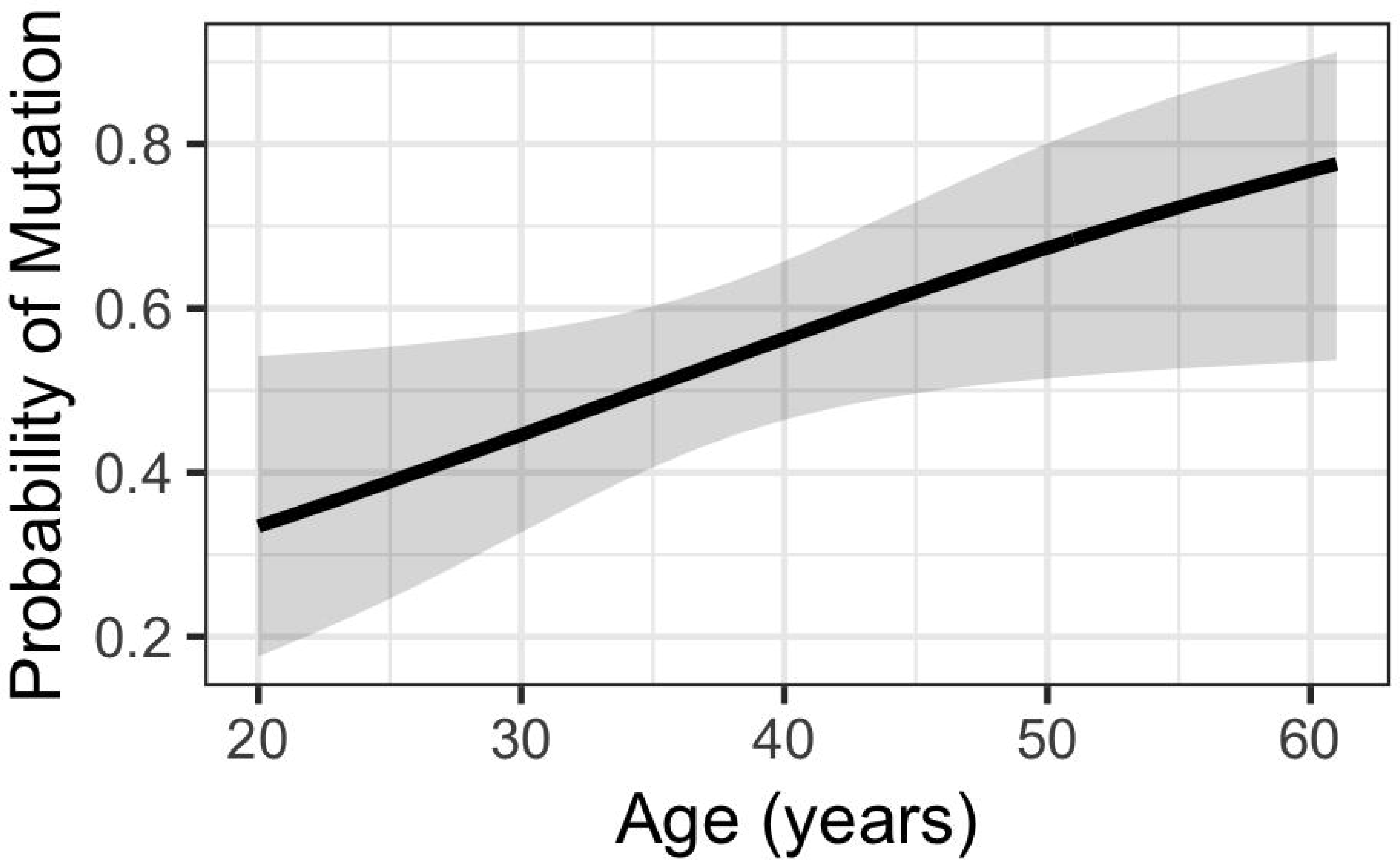
The prevalence of somatic driver mutations in the eutopic endometrium across age. According to this logistic regression model, the prevalence of mutation increases with age (OR: 1.05 (95% CI: 1.00 – 1.10), p = 0.035, Wald test).

### Comparison of Normal Endometrium Findings with Endometrial Cancer

We compared our data with endometrial cancer data from The Cancer Genome Atlas (TCGA)[28] to better interpret mutational findings within the normal endometrium. Because our analysis is restricted to the hotspot regions in the 33 genes covered by the FIND IT^™^ assay (Supplemental Table S3), we limited our comparison to oncogenic mutations within these regions for the TCGA dataset. For our normal endometrial tissue cohort, we pooled our data from both the Hx cohort (n=25) and Bx cohort (n=85). Overall, endometrial cancer samples more frequently harbored oncogenic mutations compared to the patients examined in our study (96% vs 54%, p < 0.0001, Student’s t-test) (Figure 3A; Supplementary Table S10). Women with endometrial cancer were far more likely to harbor a somatic mutation at a given age (OR: 10.8, 95% CI: 3.83 – 33.3, p = 1.5 e-05, Wald test) (Supplementary Figure S4).

**Figure 3.**
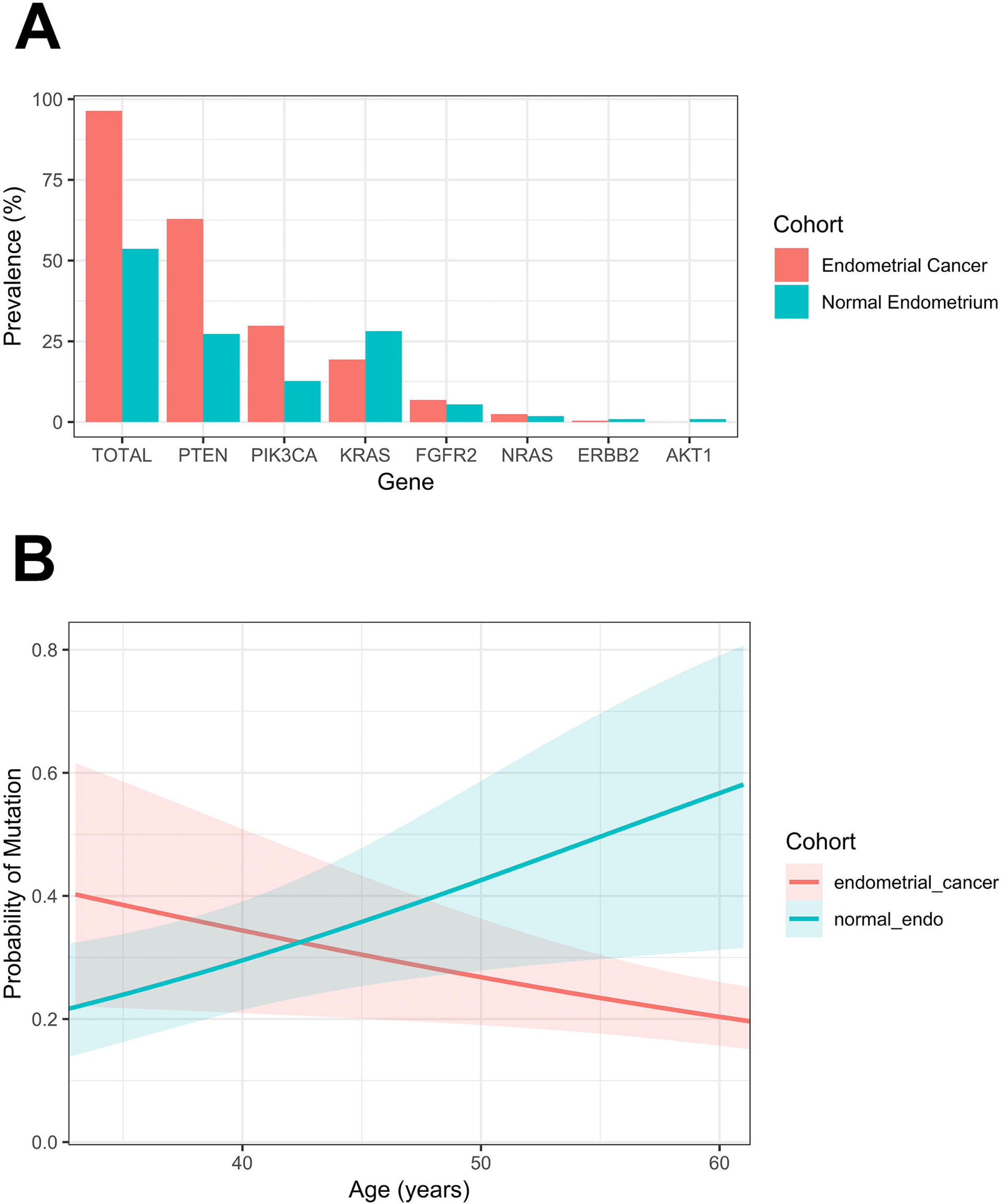
**(A)** Comparison of the prevalence of somatic mutations in endometrial cancer versus histologically normal eutopic endometrium (mutations found in endometrial cancer but were not detected in any eutopic endometrium sample are not displayed). **(B)** *KRAS* mutation prevalence across age in endometrial cancer samples versus normal endometrium samples. In normal endometrium samples, the prevalence of *KRAS* mutation increases with age (OR = 1.06, 95% CI = 1.01 – 1.11, p = 0.01953, Wald test), whereas it decreases with age in endometrial cancer samples (OR = 0.96, 95% CI = 0.94 – 0.99, p = 0.0164, Wald test).

Gene-specific analysis mutation frequencies were either indistinguishable or higher in cancer specimens, except for *KRAS* driver mutations. We found a distinct trend (p = 0.0728) suggesting a greater proportion of non-cancer patients harbored *KRAS* driver mutations in their endometrial tissue, 31 of 110 (28%), compared with endometrial cancer patients, 48 of 248 (19%) (Figure 3A; Supplementary Table S10). Indeed, in a model that incorporates age and *KRAS* mutation, whilst mutation rates in normal samples clearly increase with age (OR = 1.06, 95% CI = 1.01 – 1.11, p = 0.01953, Wald test), in the cancer samples the opposite trend is seen (OR = 0.96, 95% CI = 0.94 – 0.99, p = 0.0164, Wald test) (Figure 3B).

## Discussion

In many cancers, tumor progression is largely dependent on the accumulation of so-called “driver mutations” that confer selective growth advantages on affected subclones compared to surrounding cells[29–31]. EAOCs, in fact are characterized by mutations affecting *PIK3CA*, *PPP2R1A*, *ARID1A*, *KRAS*, *CTNNB1*, and *PTEN*[32]. Sporadic endometrial cancers also have a similar driver mutation landscape characterized by mutations affecting *PIK3CA*, *PIK3R1*, *ARID1A*, *KRAS*, *CTNNB1, PTEN*, *FGFR2,* and alterations in mismatch repair proteins[33]. The eutopic endometrium, an extremely dynamic tissue over the course of a woman’s reproductive years, is both the tissue of origin for endometrial cancer and the presumed origin of EAOCs[6–8]. Therefore, it serves as an important reference for the studies of these cancers as well as endometriosis. In the current study, we have described (with orthogonal validation) the presence of somatic, driver mutations within histologically unremarkable, eutopic endometrium of over half of women lacking evidence of gynecologic malignancy.

Our findings are consistent with oncogenic mutations existing in small clonal pockets throughout the endometrium. Amongst informative patients, oncogenic mutations were dissimilar between the anterior and posterior samplings (i.e. if a particular point mutation were found in one sampling, it was not detected in the other sampling) – these findings were confirmed by ddPCR testing of both samplings for any mutation found in either block (Table 1; Supplementary Table S5). The VAFs of reported somatic mutations were generally low (between 0.8 – 11.9% in hysterectomy or D&C/biopsy cases – see Table 1; Supplementary Table S5, S6), despite some enrichment of the glandular epithelial layer. This suggests that oncogenic mutations may be gland-level restricted, affecting only single (or a small number of) endometrial glands across a breadth of many glands from a given tissue specimen. This hypothesis seems well supported by two recent studies wherein individual endometrial glands were laser-captured from different areas within the endometrium of the same patient. Both studies suggest that different glands harbored different mutations[34,35]. These studies and our own results are also consistent with the findings by Mutter et al. (2014)[20],where PTEN-loss appears zonally in single, or small clusters of, glands (Supplementary Figure S1). Therefore, although our study represents broader sampling of the endometrium and mutations, our observations and those of others[34,35] are consistent with driver mutations, such as *KRAS* G12 mutations, occurring in small, clonal pockets throughout the uterus. Furthermore, it seems likely that acquisition of these mutations provides sufficient selective advantages to expand within an entire gland of affected endometrial cells, allowing this population to be detectable by our methods.

Somatic mutations have been previously described to accumulate with age in a variety of non-cancerous tissues[18,19,36] and thus we sought to determine the relationship between age and prevalence of somatic mutation in the normal endometrium. Overall, we found that the prevalence of oncogenic mutations in normal endometrium indeed increases by age; the risk of harboring such a mutation increases by approximately 5% per year. Moreover, the influence of age on mutation prevalence is independent of the phase of the menstrual cycle (Supplementary Figure S3). Comparing our data to the TCGA endometrial cancer dataset reveals that the prevalence of driver mutations is higher across age in endometrial cancer (or comparably as uncommon) in all genes *except* for *KRAS*. In our study, *KRAS* driver mutations occurred in over one-quarter of eutopic endometrium cases, illustrating a trend to greater prevalence in non-cancerous vs endometrial cancer samples. Although the *KRAS* mutation rates in normal endometrial samplings increased with age the opposite was seen in the TCGA cancer samples. The latter observation is likely due to distinct *KRAS* mutation rates in endometrial cancer subtypes that, themselves, have distinct age distributions[28]. Since patients in our study underwent hysterectomies, D&Cs, or endometrial biopsies for benign uterine-related pathologies or related concerns, it is plausible that *KRAS* mutations may play a role in such benign/non-cancer pathologies such as those described in arteriovenous malformations of the brain[37] or as we have previously observed in endometriosis[9,10]. It is also possible that early activation of *KRAS* induces senescence[38] in endometrial cells, thereby inhibiting progression towards malignant transformation. In contrast to our high frequency *KRAS* mutations, *CTNNB1* and *ARID1A* are commonly mutated in endometrial cancer and EAOCs[32,33] but were not observed in any of our normal endometrium specimens. This may suggest that, within the context of endometrial-derived carcinomas, such mutations are later events in tumorigenesis or that accumulation of mutations in cancer-destined precursors is etiologically distinct from age-related accumulations.

Lastly, it is important to note that many of the cancer specimens in the TCGA dataset harbor multiple driver mutations within the same tumor. Although some of our normal endometrium specimens harbored multiple driver alterations within the same block, this was relatively uncommon and it remains unclear if mutations affected overlapping populations or independent subclones. The recent study by Moore et al. found two or more driver mutations in single endometrial glands in 22% of specimens[35], suggesting an elevated probability, albeit still non-conclusive, that driver mutations may co-exist in the same non-cancerous cells.

Within the eutopic endometrium, detection of driver mutations increases with age despite the lack of remarkable histological changes. Other factors, particularly oral contraceptive use or hormonal intrauterine device use may further modulate (reduce) risk of accumulating driver mutations within endometrial tissues, such as has been observed as a decline in the frequency of PTEN-null glands in women using these contraceptive methods[21]. Because of the lack of follow-up with patients in our study, it remains unclear what role, if any, these mutations play in the eutopic endometrium. Other obvious questions include whether mutations persist, harbored in early progenitor populations, or do they disappear at each menstrual cycle re-emerging de-novo with the rapid proliferation of the endometrial lining each month[20]? Our observation of surprisingly high frequency of driver mutations in the eutopic endometrium of women without evidence of malignancy or even subtle pathology suggests that care must be taken when interpreting detected mutations in non-cancer affected individuals. Ongoing preclinical development of mutation detection assays have suggested non-cancer affected populations are unaffected by somatic driver alterations[39], many of which overlapped with our assays. However, closer inspection suggests at least some studies may not show a full picture with both sampling methodology and control population age-bias potentially contributing to the failure to detect driver mutations in non-cancer populations. The development of screening methods for occult or pre-cancerous disease will need to incorporate appropriate specimen sampling, age-appropriate control populations, and a solid understanding of normal age-related somatic mutation accumulation.

In summary, our findings highlight the complexity of the eutopic endometrium and challenge our current understanding of so-called “normal tissues”. As the mutations identified are obviously oncogenic in other tissues, these findings highlight the need to consider cell context and microenvironment along with mutation in the development of cancer.

## Acknowledgements

This research is funded by the Canadian Cancer Society (Impact grant #701603), Canadian Institutes of Health Research (Foundation grant #154290), and the Janet D. Cottrelle Foundation through the BC Cancer Foundation. The VGH & UBC Hospital Foundation and the BC Cancer Foundation provided funding to OVCARE: BC’s Ovarian Cancer Research Team. A Frederick Banting and Charles Best Canada Graduate Scholarship from the Canadian Institutes of Health Research provided support to V. Lac. The Janet D. Cottrelle Foundation Scholars fund provided support to M.S. Anglesio. The Dr. Chew Wei Memorial Professorship in Gynecologic Oncology and the Canada Research Chairs program (Research Chair in Molecular and Genomic Pathology) provided support to D.G. Huntsman. The Michael Smith Foundation for Health Research Health Professional-Investigator Award provided support to P.J. Yong.

We would also like to acknowledge the staff at the Genetic Pathology Evaluation Centre (GPEC), the Department of Anatomical Pathology at the Vancouver General Hospital in Vancouver, OVCARE’s gynaecological tissue bank, and the Calgary Laboratory Services for their assistance with specimen collection, immunohistochemical staining and optimization, and tissue sectioning.

## Statement of Author Contributions

VL, MSA, and DGH designed the study. TMN and BT-C identified specimens for the study, VL, AL, JK, TP, and MM processed samples, and performed experiments. TMN, BT-C, and MK reviewed pathology and generated/reviewed IHC data. VL and RA-H performed data analysis. AA and DC provided statistical design input and oversight. PJY and BCG provided clinical input to the study design. VL drafted the manuscript. All authors revised the manuscript and approved submission of the final version.

## Supplementary Figure Legends

**Supplementary Figure S1.**
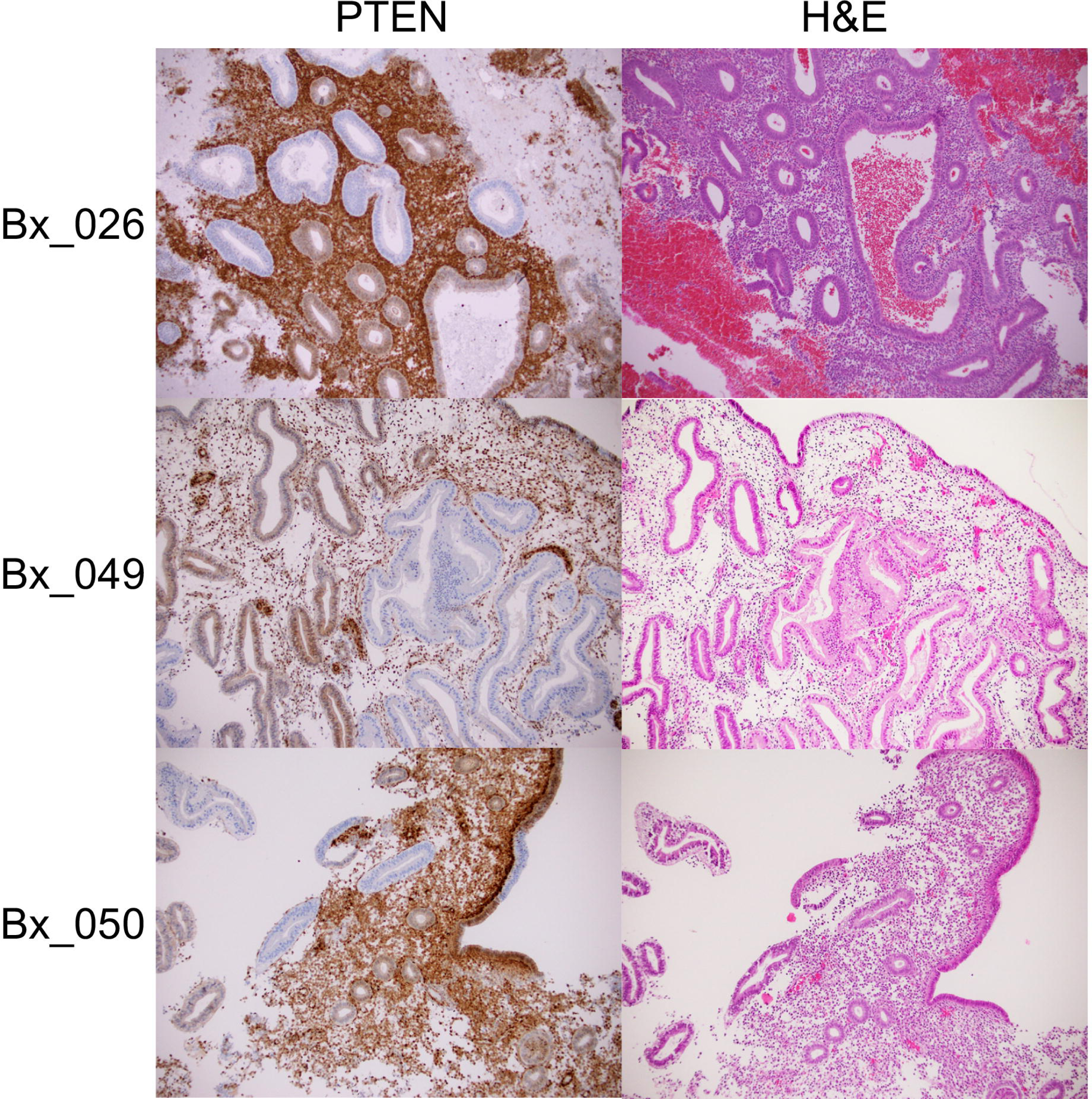
PTEN IHC studies of eutopic endometrium specimens showing zonal loss of PTEN in epithelial cells and matching H&E staining.

**Supplementary Figure S2.**
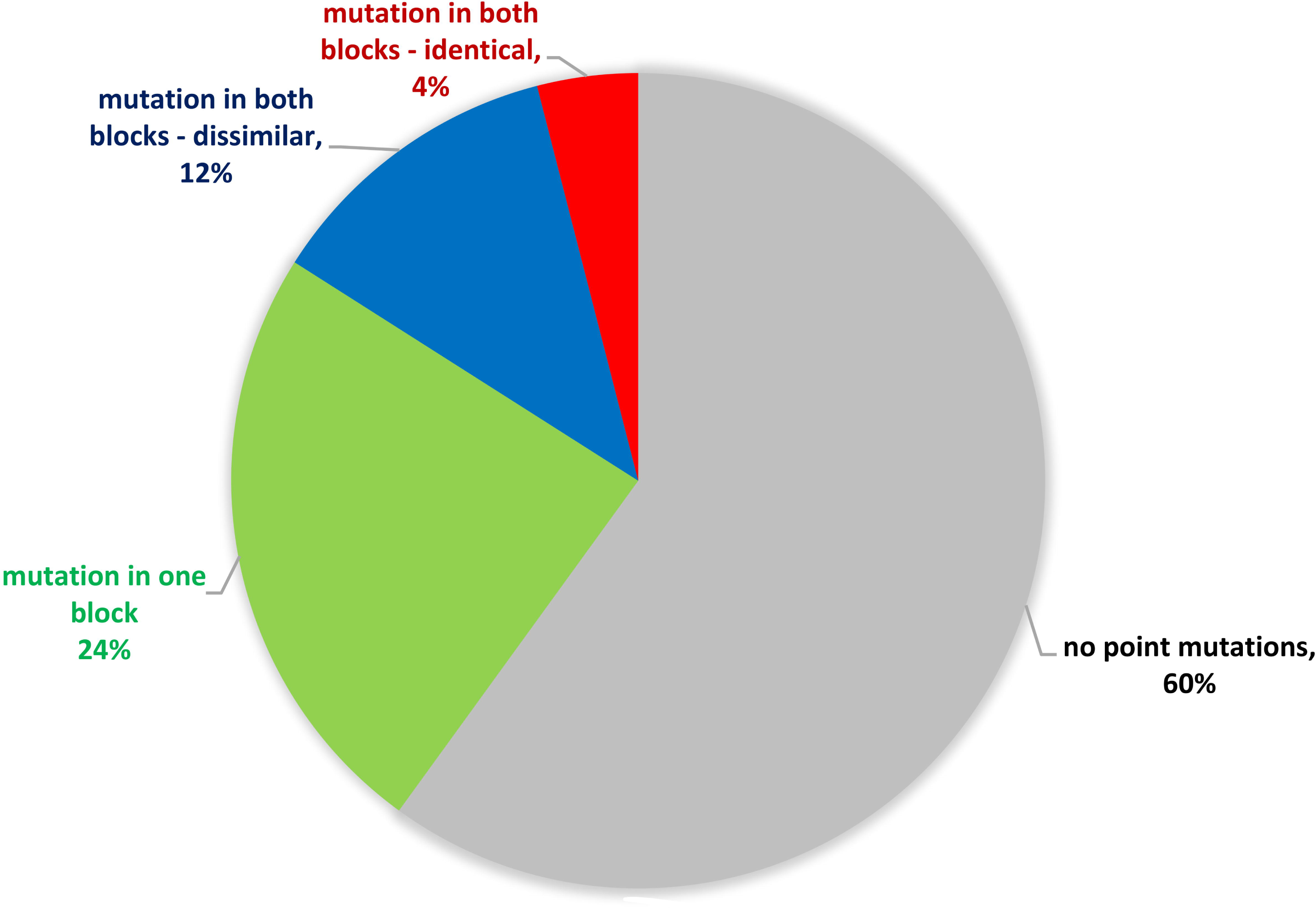
Comparison of point mutations detected in anterior and posterior samplings from patients in the Hx cohort.

**Supplementary Figure S3.**
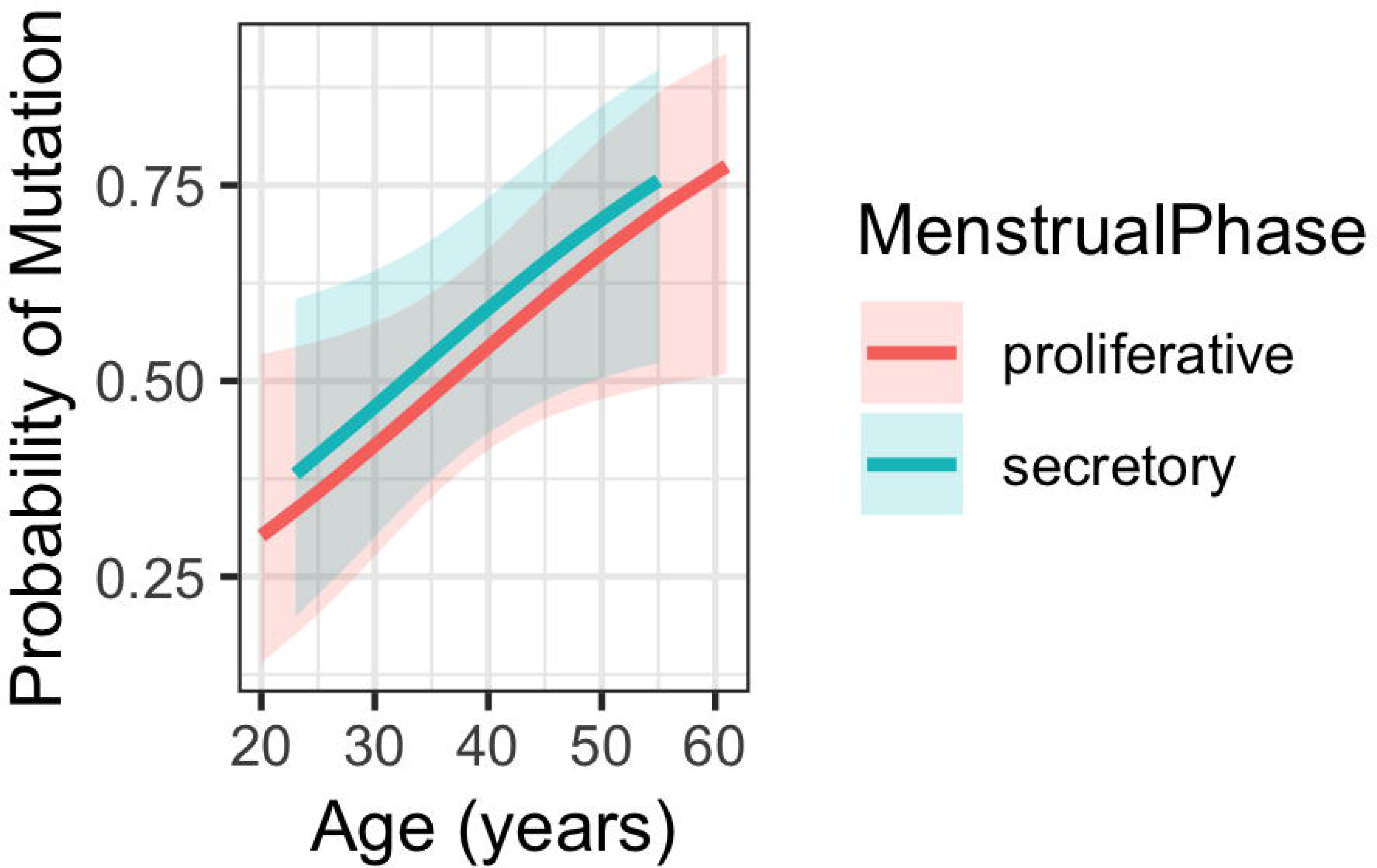
Logistic regression model of mutation prevalence across age in secretory phase versus proliferative phase endometrium

**Supplementary Figure S4.**
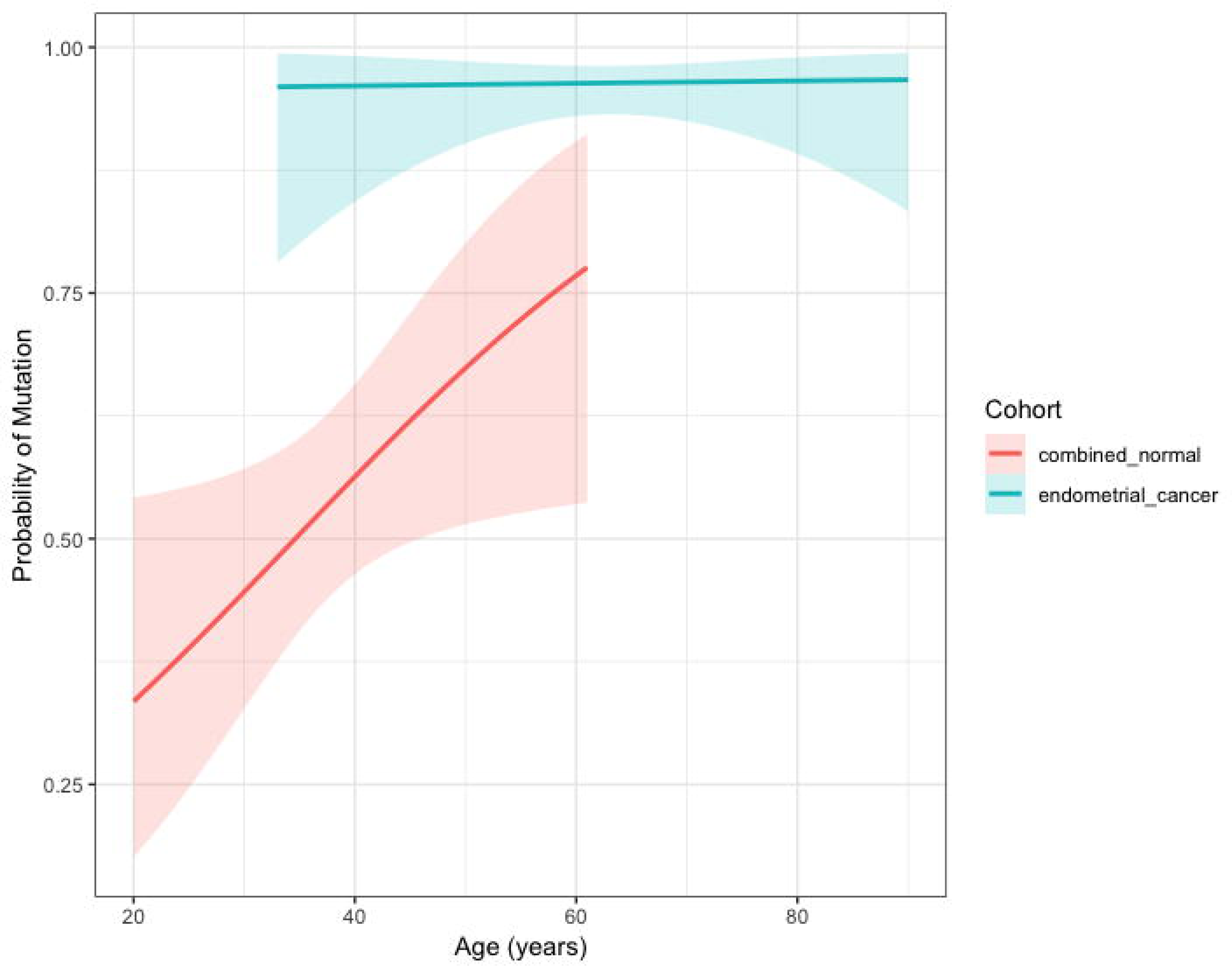
Logistic regression model of mutation prevalence across age in endometrial cancer versus normal endometrium.

